# Generation of site-specifically labelled fluorescent human XPA to investigate DNA binding dynamics during nucleotide excision repair

**DOI:** 10.1101/2023.11.23.568461

**Authors:** Sahiti Kuppa, Elliot Corless, Colleen Caldwell, Maria Spies, Edwin Antony

## Abstract

Nucleotide excision repair (NER) promotes genomic integrity by correcting bulky DNA adducts damage caused by external factors such as ultraviolet light. Defects in NER enzymes are associated with pathological conditions such as Xeroderma Pigmentosum, trichothiodystrophy, and Cockayne syndrome. A critical step in NER is the binding of the Xeroderma Pigmentosum group A protein (XPA) to the DNA adduct. To better capture the dynamics of XPA interactions with DNA during NER we have utilized the fluorescence enhancement through non-canonical amino acids (FEncAA) approach. 4-azido-*L*-phenylalanine (4AZP) was incorporated at Arg-153 in human XPA and conjugated to Cy3 using strain-promoted azide-alkyne cycloaddition. The resulting fluorescent human XPA protein (hXPA^Cy3^) shows no loss in DNA binding activity and generates a robust change in fluorescence upon binding to DNA. Here we describe methods to generate hXPA^Cy3^ and detail experimental conditions required to stably maintain the protein during biochemical and biophysical studies.

## 1. Introduction

NER is a versatile DNA repair pathway that removes a wide range of structurally unrelated adducts including N^7^-methylguanine, N^3^-methyladenine, UV-induced pyrimidine dimers, and DNA crosslinks from the genome (Li et al. 1995). Eukaryotic NER is a complex DNA repair pathway involving over a dozen protein factors that are activated upon encountering DNA damage (Lim et al. 2018). A core NER reaction can be initiated by two sub pathways: global NER (GG-NER) (Gillet and Scharer 2006) or transcription-coupled NER (TC-NER) (Hanawalt and Spivak 2008). GG-NER is a global NER pathway that can occur anywhere in the genome and detects and eliminates bulky damages in the entire genome including untranscribed regions and silent chromosomes. TC-NER, however, is specific to lesions in the transcribed strands. It operates when damage to transcribed DNA strands limit transcription. Hence, events involved in an NER pathway are often dictated by the cell cycle context. GG-NER is triggered upon DNA damage recognition initiated by the XPC-RAD23B complex, UV-DDB (UV-DNA-binding protein) (Sugasawa 2010), or DDB1-DDB2 heterodimers (XPE factors). TC-NER is activated upon damage recognition by XPC. Unlike GG-NER, TC-NER utilizes elongating Pol II to scan the transcribed strand and ‘recognize’ transcription-stalling damage. Upon recognition, the damage in both pathways is verified by the XPC-HR23B-Cen2 and TFIIH complexes. The TFIIH complex enables the downstream assembly of different NER factors such as XPA, XPB, XPC, XPD, XPG and RPA.

The resulting cascade of events can be summarized as follows: i) NER is initiated upon recruitment of xeroderma pigmentosum type C (XPC), and the transcription factor complex (TFIIH) to the damage and the DNA duplex is unwound/opened (Volker et al. 2001; Bunick et al. 2006). ii) The resulting ssDNA is stabilized through the binding of XPA and the high-affinity ssDNA binding protein replication protein A (RPA) (Matsuda et al. 1995; Li et al. 1995; Krasikova et al. 2018). Once assembled, this large complex recruits other downstream repair factors to resolve the DNA adduct (Volker et al. 2001). iii) XPF-ERCC1 and XPG, the 5′ and 3′ endonucleases respectively are next recruited to excise the bulky adduct containing DNA strand (Volker et al. 2001; Abdullah et al. 2017; Seol et al. 2018). iv) Finally, the gap is filled by DNA polymerase δ or ε in conjunction with the PCNA-RFC complex (Lim et al. 2018; Volker et al. 2001).

Here, we focus on the human XPA protein (hXPA), encoded by one of the seven XP genes (*XPA* to *XPG*) identified through genetic complementation studies of the human DNA repair disease Xeroderma Pigmentosum (Cleaver 2005). Structurally, hXPA is a 273 residue (40 kDa) protein with a partially structured Zn^2+^-containing subdomain known to coordinate DNA binding and a disordered 97 amino acid tail (Ikegami et al. 1998). hXPA interacts with almost all NER factors and plays a pivotal role in the repair process. Rad14 is the yeast homolog of hXPA and has been shown to be involved in DNA damage recognition (Koch et al. 2015). The N-terminal region of XPC directly interacts with hXPA (Bunick et al. 2006; Krasikova et al. 2010). The recruitment of the hXPA-RPA complex onto DNA is a key step in NER (Li et al. 1995). hXPA has been proposed to function as a homodimer in solution (Ikegami et al. 1998), however, recent structural studies of the NER complex show XPA as only a monomer (Kim et al. 2023; Kokic et al. 2019) during DNA damage recognition activity and a switch to the hXPA-RPA complex occurs and is proposed to promote its DNA interaction activity (Sugitani et al. 2017). The precise role of how hXPA functions in NER remains unclearly defined, thus, a detailed mechanistic characterization of the binding, dissociation, and remodeling (collectively termed “dynamics”) of hXPA on DNA in the absence and presence of RPA and other NER proteins is required. To enable such studies, we have developed a fluorescent version of hXPA that retains DNA binding activity and reports on the dynamics of hXPA-DNA interactions.

This fluorescent version of hXPA enables protein induced (or photoisomerization-related) fluorescence enhancement (PIFE) as a readout of for hXPA dynamics(Nguyen et al. 2019; Ploetz et al. 2023; Hwang and Myong 2014). For PIFE, positioning of a site-specific fluorophore is required, and the choice of position must accomplish two objectives: a) the fluorophore should produce a change in fluorescence (either an increase or decrease) upon binding of the protein to DNA, and b) the fluorescent hXPA protein should not lose activity in relation to the unlabeled protein. In terms of the structure of hXPA, the N-terminal region harbors the RPA binding motif followed by the Zn^2+^ finger region (Figure 1a) (Ikegami et al. 1998). There is one defined DNA binding domain (DBD) followed by a disordered region. We used an AlphaFold generated model (PDB: F2Z2T2; Figure 1a) and the structure of the yeast homolog (Rad14: PDB 5A39; Figure 1b) as guides for fluorophore placement (Koch et al. 2015). We show here that incorporating a fluorophore at Arg-158 in hXPA accomplishes both objectives. This residue is positioned close to the DNA binding region in the DBD and situated in a disordered loop (Figure 1b).

**Figure 1.**
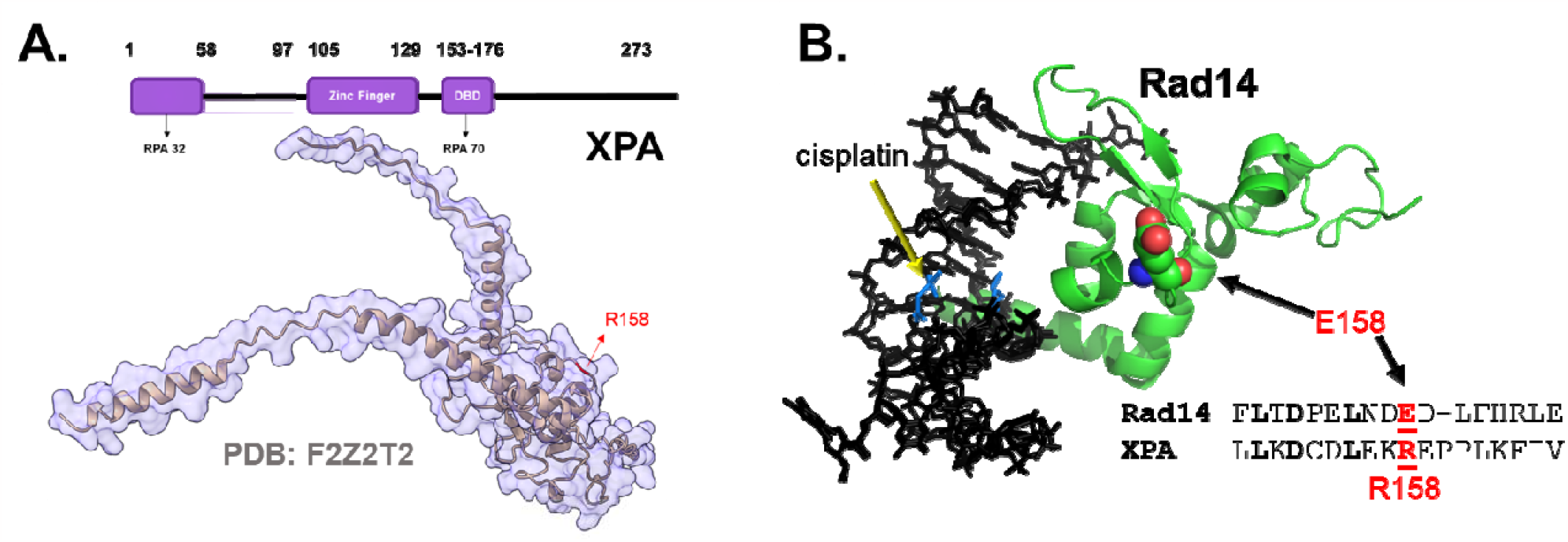
Position of 4AZP incorporation in XPA. **A)** AlphaFold model of hXPA (PDB: F2Z2T2) is shown with Arg-158 in the DNA binding domain highlighted. Arg-158 is replaced with 4AZP in hXPA^4AZP^ and tethered to a fluorophore (Cy3) in hXPA^Cy3^. **B)** Crystal structure of Rad14, the *S. cerevisiae* XPA homolog bound to DNA (PDB: 5A3D) is shown with Glu-158 highlighted. This residue is positioned close to the DNA binding region but does not make direct contact with DNA.

Since hXPA has seven Cys residues using maleimide-based fluorophore attachment is not a viable strategy. Thus, we used the non-canonical amino acid strategy to incorporate an azide handle to enable click-chemistry based fluorophore attachment (Jackson, Hammill, and Mehl 2007; Peeler and Mehl 2012). An amber suppresser codon (TAG) is engineered at the desired site in the DNA coding sequence for the hXPA protein. Recombinant protein overproduction is carried out in *E. coli* cells co-transformed with two plasmids: one carrying the hXPA open reading frame carrying the TAG substitution and a second plasmid that carries both the engineered tRNA and tRNA synthetase specific to the non-canonical amino acid of choice.

We used 4-azido-*L*-phenylalanine (4AZP) as the non-canonical amino acid and covalently attached DBCO-Cy3 using strain-promoted cyclo-addition click chemistry (Kumaresan et al. 2000; Kuppa et al. 2021; Bednar et al. 2021; Jackson, Hammill, and Mehl 2007; Morey et al. 2021). Detailed methods for generating Cy3-labeled hXPA (hXPA^Cy3^) are detailed below. hXPA^Cy3^ was purified post-labeling and the quality of the protein was assessed by SDS-PAGE (Figure 2a). While the procedure for tethering Cy3 as a fluorophore to hXPA is described here, the same approach can be used for Cy5, MB543, FITC, Atto dyes, etc. To measure activity, we compared the DNA binding activity of hXPA, hXPA carrying the 4AZP (hXPA^4AZP^), and the Cy3-tethered protein (hXPA^Cy3^). We followed the change in tryptophan (Trp) fluorescence upon DNA binding to compare the binding affinities of the three versions of hXPA (K_d_; Figure 2b). All three hXPA versions bind to DNA with comparable affinities (Figure 2b). Thus, site-specific labeling of hXPA with Cy3 at position 158 shows no observable loss in DNA binding activity (Figure. 2b). A moderate degree of Trp fluorescence quenching is observed upon DNA binding (Figure 2c), however, a robust increase in Cy3 fluorescence is observed for XPA^Cy3^ upon DNA binding resulting in a large enhancement of the signal/noise ratio compared to Trp quenching (Figure 2d). Thus, this fluorescent version of hXPA accomplishes both our initial objectives: There is no loss in activity, and the chosen position of Cy3 incorporation robustly reports on DNA interactions.

**Figure 2.**
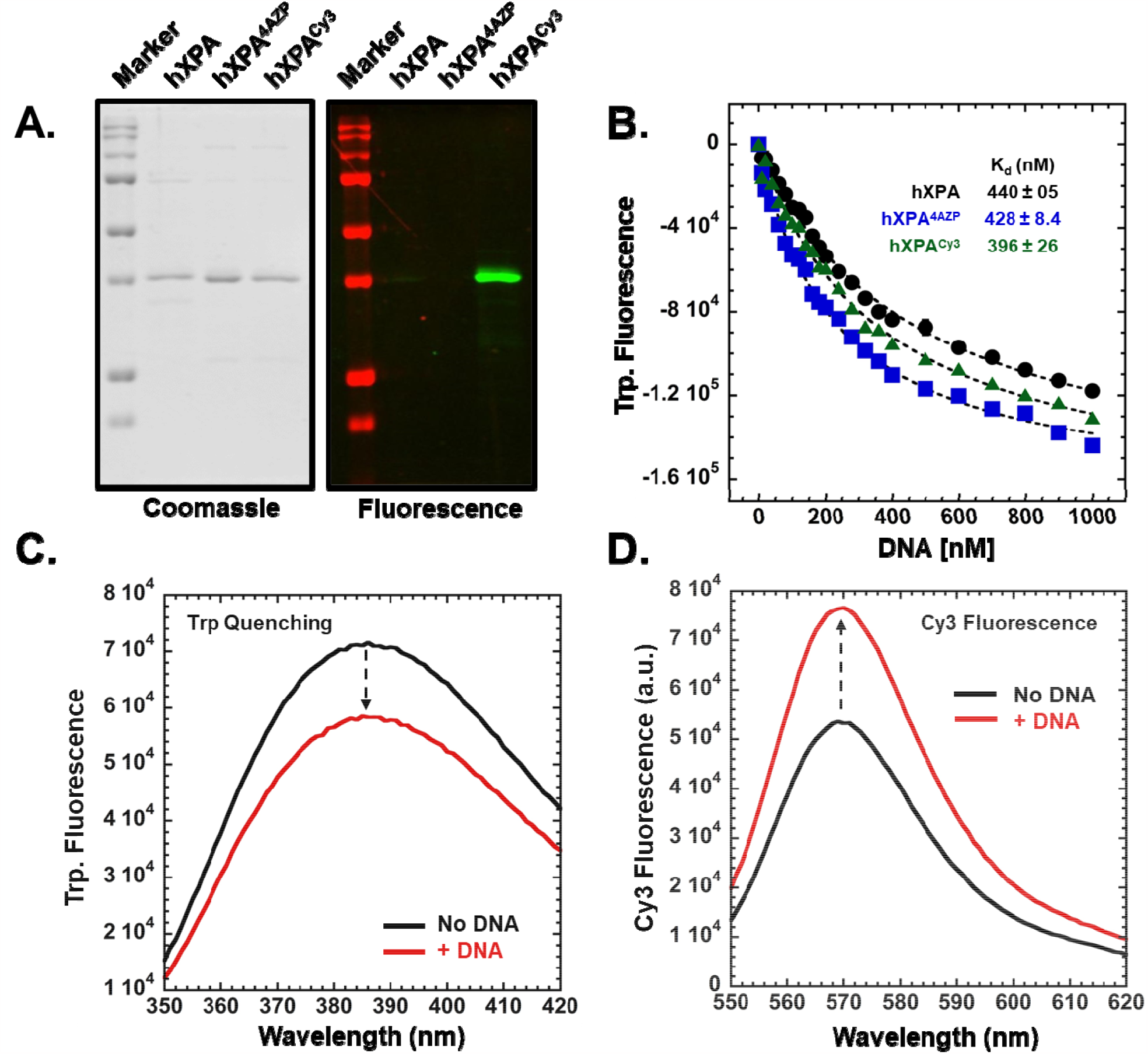
Generation of hXPA^Cy3^ and characterization of DNA binding activity. **A)** hXPA, hXPA^4AZP^, and hXPA^Cy3^ were analyzed on SDS-PAGE and subjected to imaging using Coomassie staining (left) or fluorescence imaging (right). Only hXPA^Cy3^ is observed under the fluorescence channel. **B)** DNA binding activity of hXPA, hXPA^4AZP^, and hXPA^Cy3^ were compared by monitoring the change in intrinsic Trp. fluorescence as a function of [DNA]. The DNA substrate has a 5′-(dT)_60_ nt ssDNA overhang and a 13 bp dsDNA region. 100 nM protein was used in the experiment. Data were fit to a Menten-hyperbola to obtain apparent K_d_ values for binding. For hXPA^Cy3^, **C)** a quenching in Trp. fluorescence is observed upon binding to DNA. However, **D)** an enhancement in Cy3 fluorescence is observed in the presence of DNA. These data show that hXPA^Cy3^ does not show a loss in overall DNA binding activity and the position of the Cy3 produces an excellent change in fluorescence that can be used to monitor DNA binding dynamics.

Two assays to investigate the dynamics of hXPA^Cy3^ binding to DNA are shown. The first is a stopped flow experiment where hXPA^Cy3^ is rapidly mixed with DNA and the kinetics of DNA binding are captured by following the change in Cy3 fluorescence over a short time scale providing high time resolution (Figure 3a). A single binding phase is observed which likely reflects binding and/or remodeling of the DBD of hXPA^Cy3^. The second is a single-molecule total internal reflectance fluorescence (smTIRF) microscopy experiment where DNA is tethered onto a PEG-coated glass surface and hXPA^Cy3^ is flowed onto the slide. The fluorescence traces of individual molecules are measured over time, providing high spatial resolution (Figure 3b). XPA^Cy3^ transitioning between three dynamic fluorescence states are observed (Figure 3b). These results show that the FEncAA strategy to fluorescently label hXPA is robust and provides an exciting experimental approach to investigate the DNA binding properties of hXPA.

**Figure 3.**
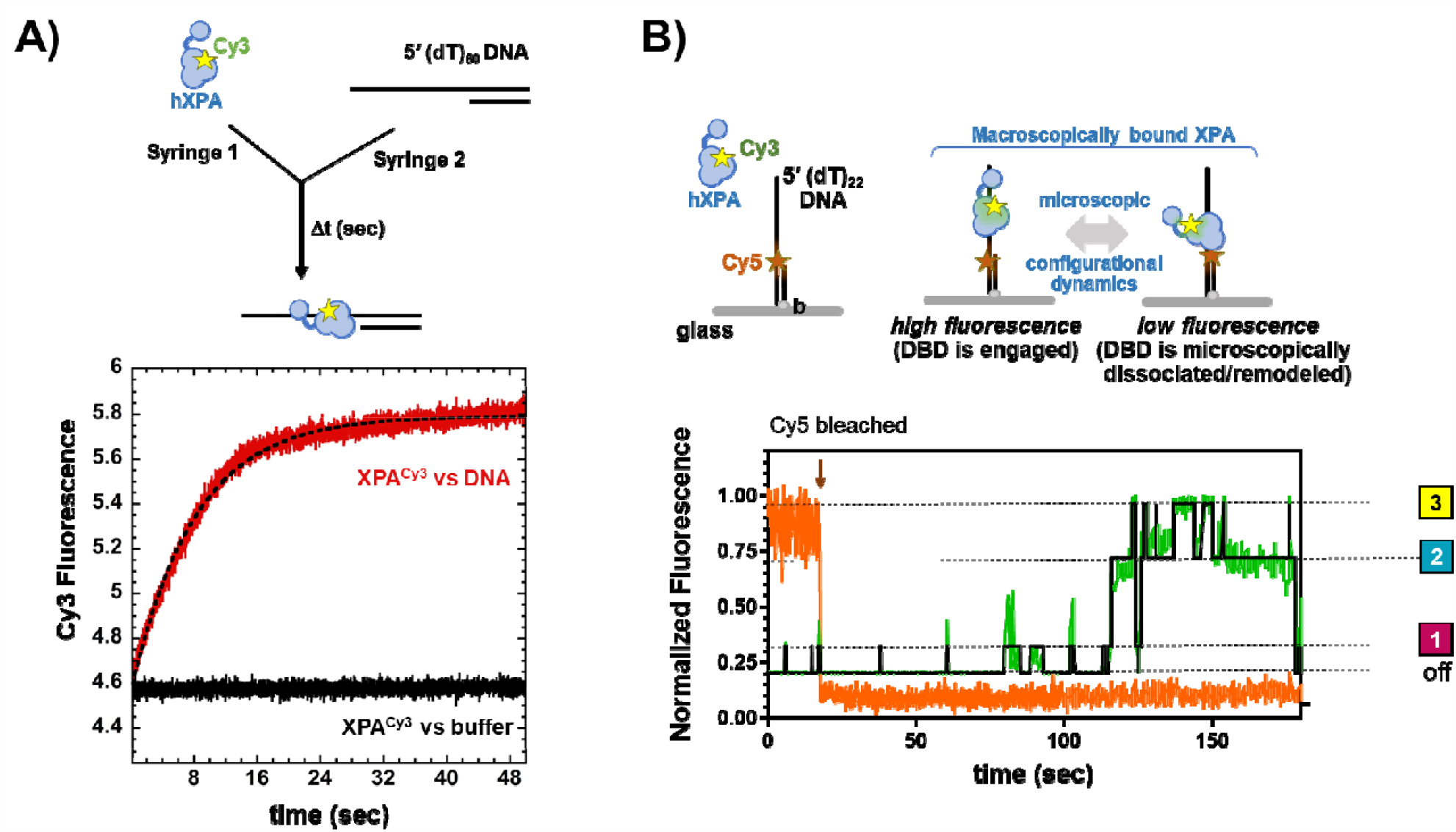
Assays to capture the DNA binding dynamics of hXPA. **A)** Stopped flow analysis of hXPA^Cy3^ binding to DNA was measured by rapidly mixing the protein and DNA (red) or buffer (black) and monitoring the change in Cy3 fluorescence. A rapid increase in Cy3 fluorescence is captured when hXPA^Cy3^ binds to DNA. The data is phenomenologically fit to a single exponential equation and yields k_obs_ = 0.12 s^-1^. **B)** Single molecule total internal reflectance fluorescence (smTIRF) microscopy of DNA binding dynamics of hXPA^Cy3^. DNA was tethered to a glass surface. Cy5 on the DNA (as pictured) was used to identify spots of binding and then photobleached. Changes in Cy3 fluorescence were measured after flowing in hXPA^Cy3^. Three difference fluorescence states are observed likely denoting multiple configurational states of the bound XPA.

In the following sections, we describe a detailed step-by-step procedure to recombinantly overproduce and purify hXPA carrying 4AZP in *E. coli* and attachment of fluorophores using strain promoted click chemistry to investigate the DNA binding dynamics of proteins involved in DNA repair.

## 2. Materials

### 2.1. Buffer Solutions

All solutions were prepared with reagent-grade chemicals. The protease inhibitor cocktail (PIC) used in our protein purifications was purchased from Millipore Sigma (catalog #P2714) and prepared as per the manufacturer’s instructions to generate a 1000X stock.

a. Cell resuspension buffer: 20mM Tris-HCl pH 7.5, 500mM NaCl, 0.5mM ZnCl_2_, 10mM imidazole, 10% glycerol (V/V), and 1.5X PIC
b. Ni^2+^-NTA Wash Buffer: 20mM Tris-HCl pH 7.5, 500mM NaCl, 30mM imidazole, 10% glycerol (V/V), and 3X PIC
c. Ni^2+^-NTA Elution Buffer 1: 20mM Tris-HCl pH 7.5, 500mM NaCl, 50mM imidazole, 10% glycerol (V/V), and 3X PIC
d. Ni^2+^-NTA Elution Buffer 2: 20mM Tris-HCl pH 7.5, 500mM NaCl, 100mM imidazole, 10% glycerol (V/V), and 3X PIC
e. Ni^2+^-NTA Elution Buffer 3: 20mM Tris-HCl pH 7.5, 500mM NaCl, 300mM imidazole, 10% glycerol (V/V), and 3X PIC
f. Ni^2+^-NTA Elution Buffer 4: 20mM Tris-HCl pH 7.5, 500mM NaCl, 500mM imidazole, 10% glycerol (V/V), and 3X PIC
g. XPA Gel Filtration Buffer: 20mM Tris-HCl pH 7.5, 500mM NaCl, 10% glycerol (V/V), and 3X PIC
h. XPA Labeling Buffer: 20mM Tris-HCl pH 7.5, 250mM NaCl, 10% glycerol (V/V), and 3X PIC
i. XPA Storage Buffer: 20mM Tris-HCl pH7.5, 500mM NaCl, 40% glycerol (V/V), and 3X PIC

l. XPA Reaction Buffer: 30mM HEPES pH 7.8, 100mM KCl, 5mM MgCl_2_, 1mM β-mercaptoethanol, and 6% glycerol (V/V).

### 2.2. Plasmids for XPA recombinant overproduction and 4AZP incorporation

The plasmid for human XPA overproduction (pHis-V5Tev) was a kind gift from Dr. Carlos Peneda (University of St. Andrews). This plasmid was modified using Q5-site directed mutagenesis (New England Biolabs, Ipswich, MA) to first convert the amber stop codon (TAG) at the end of the open-reading frame to an ochre stop codon (TAA). Next, a TAG stop codon was introduced at a position corresponding to Arg-158 in the coding region. This EA-pEV5-His-TEV-hXPA-R158TAG plasmid was used for all further studies. The TAG stop codon dictates the site for incorporation of *p*-azido-*L*-phenylalanine (4AZP). The pDule2-PCNF plasmid carrying the tRNA and tRNA synthetase specific for 4AZP incorporation was a kind gift from Dr. Ryan Mehl (Oregon State University).

## 3. Methods

### 3.1 Overproduction and purification of 4AZP-incorporated XPA

#### Day 1

Co-transform 600 ng each of the pEV5-His-TEV-hXPA-R158TAG plasmid and pDule2-PCNF plasmids into BL21 (DE3) C+ competent cells (Novagen Inc.). Plate the cells on LB-agar with kanamycin (50 µg/ml) and spectinomycin (50 µg/ml). Incubate the plates at 37°C for 16-20 hours. ∼5-20 individual colonies should be expected.

#### Day 2: Overproduction of XPA with 4AZP

1. Add kanamycin (50 µg/ml) and spectinomycin (50 µg/ml) to sterile 50 ml Luria Broth.
2. Resuspend a colony from the co-transformants on plate into the 100 ml media and incubate at 37°C for 16 – 20 hours with shaking at 250 rpm.

#### Day 3: Growth of the XPA culture

1. After the overnight incubation, add 15ml of the overnight culture to 1L of sterile Luria Broth and start shaking the cultures at 250 rpm at 37°C. Optimal incorporation of 4AZP in hXPA is achieved when the protein is overproduced using LB media or minimal media. However, in our hands, 4AZP-carrying hXPA was produced at higher yields in LB media. If this methodology is adopted for use with other proteins, we recommend testing multiple growth media to achieve optimal yields.
2. When the cultures reach OD_600_ ∼ 1.8 to 2 place the flasks on ice for about 30 mins and allow them to cool down before inducing protein overproduction. Collect and pellet 1ml of the culture at this point and store it as ‘uninduced’.
3. Add 50mM ZnCl_2_ along with 0.5M IPTG and 1mM 4AZP. To make a fresh suspension of 4AZP (enough for addition to 1L media) weigh out 206 mg of 4AZP and resuspend in 3 ml of 0.5N NaOH by vortexing. To the dissolved 4AZP, add 3ml of sterile H_2_O. Add the entire 6 ml 4AZP solution to the 1L culture. At this point, collect 5ml of the culture and incubate it in a 15ml tube along with the other flasks. This serves as a control sample for AZP incorporation into XPA. i.e., ‘Induced without AZP’ (Figure 4a).
4. Continue incubating the culture at 16°C with vigorous shaking at 250 rpm for an additional 16-18 hours.
5. All further steps are carried out at 4ºC. Harvest the cells by centrifugation at 5000 rpm for 30 min. Decant the supernatant and re-suspend the cell pellet in 25ml of pre-chilled cell resuspension buffer. The cells can be stored at -80°C for lysis at a later date or used immediately to proceed with cell lysis and protein purification (detailed below). Note: The choice of growth media and conditions for cell growth and induction are crucial for final yields of hXPA. We observed robust hXPA expression and 4AZP incorporation in LB media compared to minimal media. Similarly, overnight induction at 16ºC produced the best yields of hXPA^4AZP^ (Figure 4a). If this methodology is adopted for use with other proteins, we recommend testing multiple growth media and induction conditions to achieve optimal results.

**Figure 4.**
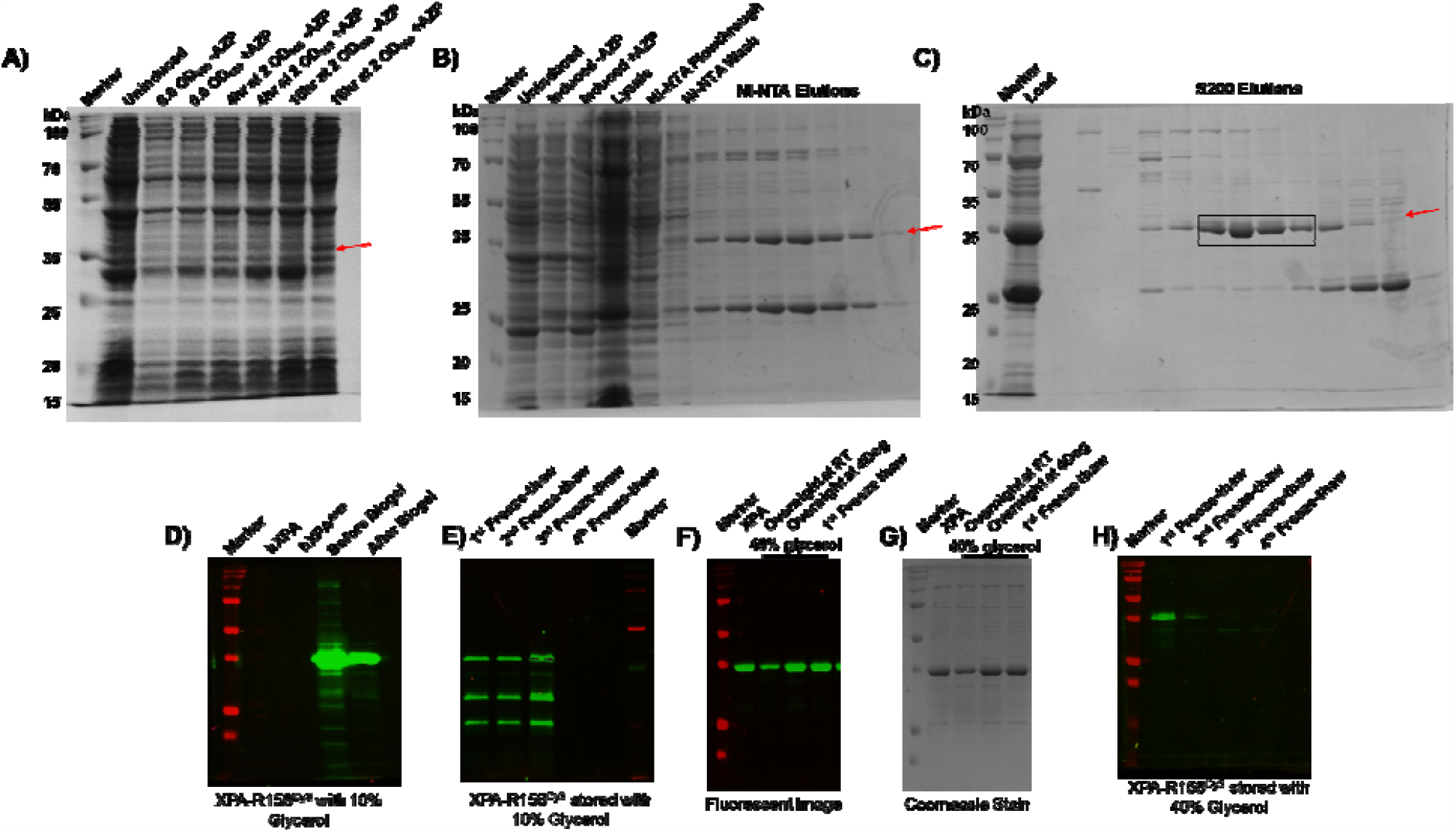
Overproduction, purification, and labeling of hXPA. **A)** SDS-PAGE analysis of uninduced and IPTG induced cells and induction carried out at varying conditions as denoted. Ideal overproduction of hXPA^4AZP^ is observed after overnight incubation at 16°C. SDS-PAGE analysis of protein samples from various fractionation steps during **B)** affinity purification on Ni^2+^-NTA and **C)** size exclusion chromatography. **D)** hXPA^Cy3^ samples before and after separation on Biogel-P4 are shown. **E)** Integrity of flash frozen hXPA^Cy3^ samples were assessed after multiple cycles of free-thaw cycles. hXPA^Cy3^ tolerates one freeze-thaw cycle when stored in 40% glycerol. **F)** hXPA^Cy3^ protein degrades when stored in buffer containing 10% glycerol. **G)** Fluorescence and **H)** Coomassie imaging of SDS-PAGE gels showing hXPA^Cy3^ stored in 40% glycerol. No degradation is observed when protein is left overnight at room temperature or at 4°C.

### Purification of XPA^4AZP^

1. Add 2ml of freshly prepared 1M phenylmethylsulfonyl fluoride (PMSF) to the resuspended cells.
2. Lyse the cells with 0.1mg/ml lysozyme and incubate at 4°C for 30 min with gentle stirring followed by sonication. Settings we use include sonicating the sample while on ice. 1 min pulse at 50 % amplitude (without stirring), stir the cells for 1 min to better mix the sample. Stop stirring and leave on ice for 1 min. Repeat 1 min pulse at 50 % amplitude without stirring.
3. Clarify the lysate by centrifugation at 17,000 rpm for 1 hour and separate the supernatant.
4. Prepare 10 ml resuspended Ni^2+^-NTA resin by equilibration with 50 mL (5 column volumes or “CV”s) of cell resuspension buffer. Allow to settle or gently spin (∼500xG) and decant most (∼40mL) of the buffer.
5. Batch bind the clarified lysate onto equilibrated Ni^2+^-NTA resin overnight at 4°C with gentle rocking.
6. Transfer the resin to an empty gravity-flow glass column and allow the lysate to flow through. The hXPA should be bound to the resin.
7. Wash the column to remove non-specifically bound proteins by sequentially washing the column with Ni^2+^-NTA Wash Buffer (100 ml).
8. Elute bound hXPA^4AZP^ with Ni^2+^-NTA Elution Buffers (20 ml) and collect the eluates.
9. Analyze the eluates on a 10% SDS-PAGE gel and pool fractions containing XPA^4AZP^ (Figure 4b).
10. Concentrate the pooled XPA^4AZP^ to ∼5 ml using a spin concentrator (10 kDa cut-off) and fractionate it over a HiLoad 16/600 Superdex 200pg size exclusion column (S200 column) using XPA Storage buffer. XPA Storage buffer is filtered using a 0.45 μm filter before use with the S200 column.
11. The largest peak eluting off the S200 column should be hXPA^4AZP^ (Figure 4c).
12. Pool the fractions containing hXPA^4AZP^ and concentrate as required using a spin concentrator (10 kDa cut-off). Measure the concentration of hXPA^4AZP^ using extinction coefficient (ε_280_) 28795 M^-1^cm^-1^ or Bradford Assay.

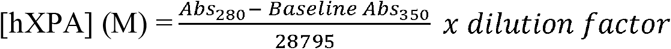
13. Flash freeze and store hXPA^4AZP^ as small aliquots in liquid nitrogen, then store at -80 °C, or proceed to labeling reactions (below).
14. Site-specific incorporation of 4AZP can be checked using mass spectrometry (if desired).
15. We noticed that hXPA is prone to degradation upon multiple (2) freeze thaws and buffer conditions needed optimization to prevent this issue. First, we noticed that hXPA^Cy3^ eluted as a clean fraction after separation on Biogel-P4 (Figure 4d). However, hXPA^Cy3^ stored with 10% glycerol degraded during storage (Figure 4e). Adjusting the final concentration of glycerol to 40% for long-term storage significantly improved the stability of hXPA^Cy3^ (Figure 4f & g). Under these conditions, the protein is stable overnight at RT or at 4ºC. The flash frozen protein is stable for one freeze-thaw cycle (Figure 4h). But, even with higher glycerol containing storage buffer, more than one freeze-thaw cycle should be avoided. We highly recommend tracking the condition of hXPA^Cy3^ before every experiment.
16. Final yields of hXPA^4AZP^ are ∼ 0.2 - 0.3 mg/ml and 1.5 mg total protein per L of growth with 4AZP incorporation.

### Labeling of XPA^4AZP^ P with click-chemistry based fluorophores

*Choice of fluorophores and click chemistry options*. 4AZP can be conjugated to an alkyne group carrying fluorophores using either Cu-based or Cu-free click chemistry. We have successfully applied the procedure below for use with the following Cu-free fluorophores: DBCO-Cy3, DBCO-Cy5, and DBCO-MB543, and is likely applicable to other DBCO-derivatives. But the reaction conditions must be systematically determined. In this case, these fluorophores were sourced from Click Chemistry Tools Inc. (Scottsdale, AZ).

1. The fluorophores were resuspended as per the manufacturer’s instructions. Typically, we use dimethyl formamide (DMF) to resuspend the dye. Measure the concentration of the resuspended dye. Measure the concentration of the fluorescent dye using the following formula:

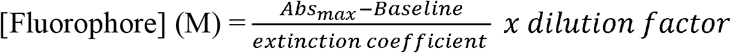
2. Click chemistry labeling reactions were performed in 3 ml – 5 ml total volume. Reactions contained ∼2 – 4 µM hXPA^4AZP^ and a 2-fold molar excess of the DBCO-fluorophore.
3. Labeling reactions were performed at 4°C for 2 hours using a gentle rocker and the reaction tubes were covered using tin foil.
4. Next, the reaction was loaded onto a Biogel-P4 column (40cm x 2cm) to separate labeled hXPA^4AZP^ from excess dye. Equilibrate the Biogel-P4 column with hXPA Storage buffer, load the reaction, and resolve the labeled-hXPA form the excess dye. We typically run our Biogel-P4 column at 0.1 ml/min on the FPLC. The fluorescently labeled hXPA will elute out as the leading band and trailing diffused band will be the excess dye.
5. Pool fractions containing fluorescent-hXPA. Add glycerol to achieve an adjusted final concentration of 40% (v/v). Please see note about hXPA^Cy3^ stability noted above.
6. Measure the concentration and labeling efficiency, and flash freeze the protein. Store the samples at -80 °C. Fluorescent versions of hXPA are stable for up to 2 months at -80°C. Also avoid multiple freeze-thaw cycles as hXPA^Cy3^ samples will degrade.
7. The labeling efficiency for hXPA^4AZP^ with DBCO-Cy3 is usually about ∼ 60 %.

### Measure the efficiency of labeling

1. To test dye conjugation, analyze unlabeled and labeled hXPA on a 10% SDS-PAGE gel. Image the gel using a fluorescence imager and with laser settings appropriate for the fluorescent dye. For Cy3, we use the 515-545 nm channel on an iBright imager (Thermo Fisher Scientific). Only the fluorescent hXPA band should be visualized in the fluorescent image. Next, Coomassie stain the same gel and you should be able to visualize hXPA in both the unlabeled and labeled samples.
2. Measure the concentration of XPA and the fluorophore in the labeled protein sample using a UV-Vis spectrophotometer. The respective extinction coefficients of the protein and the fluorophore should be used to determine the concentration and the labeling ratios. Described below.
3. Correction Factor: Please note that it will be useful to measure the baseline absorbance spectra of the excess dye eluted from the Biogel-P4 separation. This will provide the absorbance contribution of the fluorophore at 280 nm and this value should be used to correct for hXPA concentration measured at 280 nm for the fluorescent protein.

### Corrected Measurement of Protein Concentration using Absorbance

1. Perform an absorbance scan of the fluorescent hXPA sample from 240nm to 700nm (or up to the wavelength of the desired fluorophore used).
2. Perform an absorbance scan of the free dye eluted from the Biogel-P4 column. Scan from 240nm to 700nm (or up to the wavelength of the desired fluorophore used). Pick the absorbance maxima at 280nm and at the optimal maxima for the fluorophore used. Then, generate a ‘Correction Factor’ using these two values and the following formula.

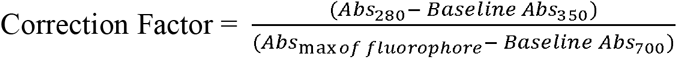
3. Next, calculate the total concentration of hXPA from the absorbance maxima at 280nm using this formula: 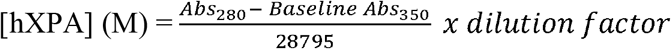
4. Next, apply the correction factor measured above to correct for fluorophore contributions at 280nm. You can use this general formula to achieve this:

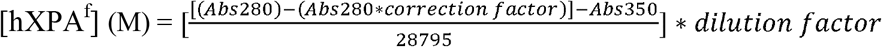
5. Measure the concentration of the attached fluorophore using the corresponding value at the absorption maxima and the appropriate extinction coefficient for the fluorophore used.

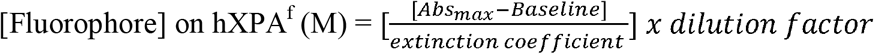
6. The ratio of the fluorophore versus protein concentration yields the labeling efficiency.

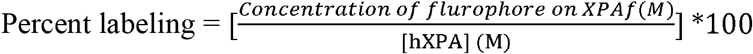

### Testing the DNA binding activity of labeled hXPA

#### Steady-state changes in DNA binding induced hXPA^Cy3^ fluorescence

Changes in intrinsic tryptophan fluorescence were used to capture hXPA^WT^, hXPA^AZP^ or hXPA^f^ binding to DNA in reaction buffer (Figure 2). Fluorescence spectra were obtained using a PTI QM40 instrument (Horiba Scientific, Edison, NJ, USA). 100nM hXPA was mixed with 0-1 μM DNA (a 13 base pair dsDNA substrate with a 60 nt 5′ ssDNA overhang) in reaction buffer (30mM HEPES, pH 7.8, 100mM KCl, 5mM MgCl_2_, 1mM β-mercaptoethanol and 6% (v/v) glycerol) and added to a 2 ml quartz cuvette. The change in Trp fluorescence was monitored by exciting the sample at 290nm and measuring emission at 325nm. All experiments were performed at 25ºC. Each data point data was collected as an average of 10 repeats. The experiment was repeated n=3. The data were phenomenologically fit to single binding site model in Prism to obtain apparent dissociation constants. Similar experiments were also performed to monitor the changes in hXPA^Cy3^ fluorescence. Here, samples were excited at 535 nm and emission scans were recorded between 550 – 620 nm.

#### Pre-steady state changes in DNA binding induced hXPA^Cy3^ fluorescence

Stopped flow experiments were performed using an Applied Photophysics SX20 instrument (Surrey, UK). 100nM hXPA^Cy3^ from one syringe was mixed with 500nM DNA (same substrate as described above) from the second syringe. Experiments were performed in reaction buffer. Samples were excited at 515nm and Cy3 fluorescence was monitored using a 530 nm cut-off long pass filter. Seven individual reactions were averaged and the change in Cy3 fluorescence was plotted as a function of time. The traces were fit to a single exponential equation to generate an observed rate of binding and signal change.

#### smTIRF analysis of hXPA^Cy3^ DNA binding dynamics

Analysis of the DNA binding and/or configurational dynamics of hXPA can be performed on a variety of TIRFM configurations. Here, we used a bespoke prism based TIRF microscope built around an Olympus IX71 microscope frame (see Bain *et. al*. for details; (Bain et al. 2018)). The Cy3 fluorescence was excited using 532nm (Coherent, Compass 215M-50) laser, and Cy5 fluorescence was excited using 641nm (Coherent, Cube 1150205 / AD) lasers. The two laser beams were combined using a polarizing beam splitting cube (CVI Melles Griot, PBSH-450-700-050) and directed to the microscope objective at a 30° angle. TIR was achieved via a UV fused silica pellin-broca prism (Eksma Optics, 325-1206) and uncoated N-BK7 plano-convex lens (Thorlabs, LA1213). Fluorescence emitted by the Cy3 and Cy5 dyes was collected using a 60X, NA 1.20 water immersion objective (Olympus, UPLSAPO60XW), and spurious fluorescent signal was removed with a dual bandpass filter (Semrock, FF01-577/690-25). Cy3 and Cy5 emissions were separated using a dual-view (Photometrics, DV2) housing a 650nm longpass filter (Chroma Technology Corp., T650lpxr). Fluorescent images are collected using an Andor iXon 897 EMCCD (Oxford Instruments).

The biotinylated, Cy5-labeled DNA substrates (Figure 3B) were prepared by mixing of the two oligos at 1μM concentrations, heating to 95°C and slowly cooling to room temperature to facilitated annealing. Quartz slides were passivated and PEGylated prior as described in detail (Bain et al. 2018). The flow cell was treated with 0.2 mg/ml NeutrAvidin. Excess neutravidin was removed with 1mL of wash buffer (25 mM Tris-HCl pH 7.0, 140 mM KCl). The flow cell is incubated with 100pM of biotinylated partial duplex DNA substrate for 3 minutes at which time excess DNA is removed with wash buffer. Cy3-labeled hXPA was then flowed into the flow cell chamber in imaging buffer (25 mM Tris-HCl pH 7.0, 140 mM KCl, 10 mM MgCl2, 1 mg/ml BSA, 1 mM DTT, 0.8% w/v D-glucose, 12 μM glucose oxidase, 0.04 mg/ml catalase, 24 mM TROLOX). Dual excitation Cy3/Cy5 movies were then collected at 100 ms time resolution (gain of 290, background set to 400, and correction set to 1200) using custom software, single.exe, (generously provided by the Taekjip Ha Lab, JHU) with 532 nm and 642 nm laser power set to 45 mW. Movies were collected for a total of 1800 frames (180 s). Fluorescent trajectory extraction was carried out using an IDL script (generously provided by the Taekjip Ha Lab, JHU). Individual trajectories were then imported into hFRET (Hon and Gonzalez 2019) and fit to 2-6 states of fluorescent intensity. The best fit was determined from the lower bound comparison of the tested models. Four states, one unbound and three bound states were selected. Please also refer to Pokhrel et. al. for additional single molecule approaches to capture DNA binding dynamics (Pokhrel et al. 2019).

## Acknowledgements

This work was supported by MIRA grants from the National Institutes of Health and NIGMS R35GM149320 to E.A. and R35GM131704 to M.S. and the National Cancer Institute F99 CA274696 to S.K. Analytical ultracentrifugation experiments to characterize XPA were supported by a grant from the NIH office of the director S10 OD030343

## References

Abdullah, U. B., J. F. McGouran, S. Brolih, D. Ptchelkine, A. H. El-Sagheer, T. Brown, and P. J. McHugh. 2017. ‘RPA activates the XPF-ERCC1 endonuclease to initiate processing of DNA interstrand crosslinks’, EMBO J, 36: 2047–60.

Bain, F. E., L. A. Fischer, R. Chen, and M. S. Wold. 2018. ‘Single-Molecule Analysis of Replication Protein A-DNA Interactions’, Methods Enzymol, 600: 439–61.

Bednar, R. M., S. Jana, S. Kuppa, R. Franklin, J. Beckman, E. Antony, R. B. Cooley, and R. A. Mehl. 2021. ‘Genetic Incorporation of Two Mutually Orthogonal Bioorthogonal Amino Acids That Enable Efficient Protein Dual-Labeling in Cells’, ACS Chem Biol, 16: 2612–22.

Bunick, C. G., M. R. Miller, B. E. Fuller, E. Fanning, and W. J. Chazin. 2006. ‘Biochemical and structural domain analysis of xeroderma pigmentosum complementation group C protein’, Biochemistry, 45: 14965–79.

Cleaver, J. E. 2005. ‘Cancer in xeroderma pigmentosum and related disorders of DNA repair’, Nat Rev Cancer, 5: 564–73.

Gillet, L. C., and O. D. Scharer. 2006. ‘Molecular mechanisms of mammalian global genome nucleotide excision repair’, Chem Rev, 106: 253–76.

Hanawalt, P. C., and G. Spivak. 2008. ‘Transcription-coupled DNA repair: two decades of progress and surprises’, Nat Rev Mol Cell Biol, 9: 958–70.

Hon, J., and R. L. Gonzalez, Jr. 2019. ‘Bayesian-Estimated Hierarchical HMMs Enable Robust Analysis of Single-Molecule Kinetic Heterogeneity’, Biophys J, 116: 1790–802.

Hwang, H., and S. Myong. 2014. ‘Protein induced fluorescence enhancement (PIFE) for probing protein-nucleic acid interactions’, Chem Soc Rev, 43: 1221–9.

Ikegami, T., I. Kuraoka, M. Saijo, N. Kodo, Y. Kyogoku, K. Morikawa, K. Tanaka, and M. Shirakawa. 1998. ‘Solution structure of the DNA- and RPA-binding domain of the human repair factor XPA’, Nat Struct Biol, 5: 701–6.

Jackson, J. C., J. T. Hammill, and R. A. Mehl. 2007. ‘Site-specific incorporation of a (19)F-amino acid into proteins as an NMR probe for characterizing protein structure and reactivity’, J Am Chem Soc, 129: 1160–6.

Kim, J., C. L. Li, X. Chen, Y. Cui, F. M. Golebiowski, H. Wang, F. Hanaoka, K. Sugasawa, and W. Yang. 2023. ‘Lesion recognition by XPC, TFIIH and XPA in DNA excision repair’, Nature, 617: 170–75.

Koch, S. C., J. Kuper, K. L. Gasteiger, N. Simon, R. Strasser, D. Eisen, S. Geiger, S. Schneider, C. Kisker, and T. Carell. 2015. ‘Structural insights into the recognition of cisplatin and AAF-dG lesion by Rad14 (XPA)’, Proc Natl Acad Sci U S A, 112: 8272–7.

Kokic, G., A. Chernev, D. Tegunov, C. Dienemann, H. Urlaub, and P. Cramer. 2019. ‘Structural basis of TFIIH activation for nucleotide excision repair’, Nat Commun, 10: 2885.

Krasikova, Y. S., N. I. Rechkunova, E. A. Maltseva, and O. I. Lavrik. 2018. ‘RPA and XPA interaction with DNA structures mimicking intermediates of the late stages in nucleotide excision repair’, PLoS One, 13: e0190782.

Krasikova, Y. S., N. I. Rechkunova, E. A. Maltseva, I. O. Petruseva, and O. I. Lavrik. 2010. ‘Localization of xeroderma pigmentosum group A protein and replication protein A on damaged DNA in nucleotide excision repair’, Nucleic Acids Res, 38: 8083–94.

Kumaresan, S., N. Yoganandan, F. A. Pintar, D. J. Maiman, and S. Kuppa. 2000. ‘Biomechanical study of pediatric human cervical spine: a finite element approach’, J Biomech Eng, 122: 60–71.

Kuppa, S., N. Pokhrel, E. Corless, S. Origanti, and E. Antony. 2021. ‘Generation of Fluorescent Versions of Saccharomyces cerevisiae RPA to Study the Conformational Dynamics of Its ssDNA-Binding Domains’, Methods Mol Biol, 2281: 151–68.

Li, L., X. Lu, C. A. Peterson, and R. J. Legerski. 1995. ‘An interaction between the DNA repair factor XPA and replication protein A appears essential for nucleotide excision repair’, Mol Cell Biol, 15: 5396–402.

Lim, K. K., T. T. T. Nguyen, A. Y. Li, Y. P. Yeo, and E. S. Chen. 2018. ‘Histone H3 lysine 36 methyltransferase mobilizes NER factors to regulate tolerance against alkylation damage in fission yeast’, Nucleic Acids Res, 46: 5061–74.

Matsuda, T., M. Saijo, I. Kuraoka, T. Kobayashi, Y. Nakatsu, A. Nagai, T. Enjoji, C. Masutani, K. Sugasawa, F. Hanaoka, and et al. 1995. ‘DNA repair protein XPA binds replication protein A (RPA)’, J Biol Chem, 270: 4152–7.

Morey, T. M., M. A. Esmaeili, M. L. Duennwald, and R. J. Rylett. 2021. ‘SPAAC Pulse-Chase: A Novel Click Chemistry-Based Method to Determine the Half-Life of Cellular Proteins’, Front Cell Dev Biol, 9: 722560.

Nguyen, B., M. A. Ciuba, A. G. Kozlov, M. Levitus, and T. M. Lohman. 2019. ‘Protein Environment and DNA Orientation Affect Protein-Induced Cy3 Fluorescence Enhancement’, Biophys J, 117: 66–73.

Peeler, J. C., and R. A. Mehl. 2012. ‘Site-specific incorporation of unnatural amino acids as probes for protein conformational changes’, Methods Mol Biol, 794: 125–34.

Ploetz, E., B. Ambrose, A. Barth, R. Borner, F. Erichson, A. N. Kapanidis, H. D. Kim, M. Levitus, T. M. Lohman, A. Mazumder, D. S. Rueda, F. D. Steffen, T. Cordes, S. W. Magennis, and E. Lerner. 2023. ‘A new twist on PIFE: photoisomerisation-related fluorescence enhancement’, ArXiv.

Pokhrel, N., C. C. Caldwell, E. I. Corless, E. A. Tillison, J. Tibbs, N. Jocic, S. M. A. Tabei, M. S. Wold, M. Spies, and E. Antony. 2019. ‘Dynamics and selective remodeling of the DNA-binding domains of RPA’, Nat Struct Mol Biol, 26: 129–36.

Seol, J. H., C. Holland, X. Li, C. Kim, F. Li, M. Medina-Rivera, R. Eichmiller, I. F. Gallardo, I. J. Finkelstein, P. Hasty, E. Y. Shim, J. A. Surtees, and S. E. Lee. 2018. ‘Distinct roles of XPF-ERCC1 and Rad1-Rad10-Saw1 in replication-coupled and uncoupled inter-strand crosslink repair’, Nat Commun, 9: 2025.

Sugasawa, K. 2010. ‘Regulation of damage recognition in mammalian global genomic nucleotide excision repair’, Mutat Res, 685: 29–37.

Sugitani, N., M. W. Voehler, M. S. Roh, A. M. Topolska-Wos, and W. J. Chazin. 2017. ‘Analysis of DNA binding by human factor xeroderma pigmentosum complementation group A (XPA) provides insight into its interactions with nucleotide excision repair substrates’, J Biol Chem, 292: 16847–57.

Volker, M., M. J. Mone, P. Karmakar, A. van Hoffen, W. Schul, W. Vermeulen, J. H. Hoeijmakers, R. van Driel, A. A. van Zeeland, and L. H. Mullenders. 2001. ‘Sequential assembly of the nucleotide excision repair factors in vivo’, Mol Cell, 8: 213–24.

